# Automated genome mining predicts combinatorial diversity and taxonomic distribution of peptide metallophore structures

**DOI:** 10.1101/2022.12.14.519525

**Authors:** Zachary L. Reitz, Alison Butler, Marnix H. Medema

## Abstract

Microbial competition for trace metals shapes their communities and interactions with humans and plants. Many bacteria scavenge trace metals with metallophores, small molecules that chelate environmental metal ions and transport them back into the cell. Our incomplete knowledge of metallophores diversity stymies our ability to fight infectious diseases and harness beneficial microbiome interactions. The majority of known metallophores are non-ribosomal peptides (NRPs), which feature metal-chelating moieties rarely found in other classes of natural products. NRP metallophore production may be predicted by genome mining, where genomes are scanned for homologs of known biosynthetic gene clusters (BGCs). However, accurately detecting NRP metallophore biosynthesis currently requires expert manual inspection. Here, we introduce automated identification of NRP metallophore BGCs through a comprehensive detection algorithm, newly implemented in antiSMASH. Custom-designed profile hidden Markov models detect genes encoding the biosynthesis of most known NRP metallophore chelating moieties (2,3-dihydroxybenzoate, hydroxamates, salicylate, β-hydroxyamino acids, graminine, Dmaq, and the pyoverdine chromophore), achieving 97% precision and 78% recall against manual curation. We leveraged the algorithm, in combination with transporter gene detection, to detect NRP metallophore BGCs in 15,562 representative bacterial genomes and predict that 25% of all non-ribosomal peptide synthetases encode metallophore production. BiG-SCAPE clustering of 2,562 NRP metallophore BGCs revealed that significant diversity remains unexplored, including new combinations of chelating groups. Additionally, we find that Cyanobacteria are severely understudied and should be the focus of more metallophore isolation efforts. The inclusion of NRP metallophore detection in antiSMASH version 7 will aid non-expert researchers and facilitate large-scale investigations into metallophore biology.

## Introduction

Across environments, microbes compete for a scarce pool of trace metals. Many microbes scavenge metal ions with small-molecule chelators called *metallophores*, which diffuse through the environment and chelate metal ions with high affinity.^1,2^ A microbe possessing the right membrane transporters will be able to recognize and import a metallophore–metal complex, while other strains are unable to access the chelated metal ions. Thus, the metallophore excreted by one microbe can either support or inhibit growth of a neighboring strain, driving complex community dynamics in marine, freshwater, soil, and host environments.^3^ The most well studied metallophores are the Fe(III)-binding *siderophores*, which have found applications in biocontrol, bioremediation, and medicine.^4^ Two recent studies demonstrated that the disease suppression ability of a rhizosphere microbiome is strongly determined by whether or not the pathogen can use siderophores produced by the community; a microbiome can even encourage pathogen growth when a compatible siderophore is produced.^5,6^ Compared to siderophores, other metallophore classes are relatively understudied, but they likely play equally important biological roles, as exemplified by recent reports of both commensal and pathogenic bacteria relying on zincophores to effectively colonize human hosts.^7,8^

Hundreds of unique metallophore structures have been characterized, each with specific chemical properties (ex: effective pH range, hydrophobicity, and metal selectivity) and biological effects on other microbes (based on membrane transporter compatibility). Experimentally characterizing metallophores can be time-consuming and costly, and thus researchers often use genome mining to predict metallophore production in silico.^9^ Taxonomy alone is not sufficient to predict what metallophores will be produced by a microbe, as production can vary significantly even within a single species.^10^ Instead, metallophores must be predicted from each genome based on the presence of biosynthetic gene clusters (BGCs) that encode their biosynthesis. The majority of known metallophores are non-ribosomal peptides (NRPs), a broad class of natural products that also includes many antibiotics, antitumor compounds, and toxins. Specialized chelating moieties bind directly to the metal ion (in the case of siderophores, Fe^3+^), while other amino acids in the peptide chain give the metallophore the required flexibility for chelation (Figure 1A). Only a handful of chelating groups have been identified to date, and nearly all NRP metallophores contain one or more of the substructures shown in Figure 1A: 2,3-dihydroxybenzoate (catechol, 2,3-DHB), hydroxamates, salicylate, β-hydroxyaspartate (β-OHAsp), β-hydroxyhistidine (β-OHHis), graminine, Dmaq (1,1-dimethyl-3-amino-1,2,3,4-tetrahydro-7,8-dihydroxy-quinoline), and the pyoverdine chromophore. Biosynthetic pathways are known for each of the chelating groups (Figure 1B), and the presence of a chelator pathway may be used as a marker for metallophore production.

**Figure 1.**
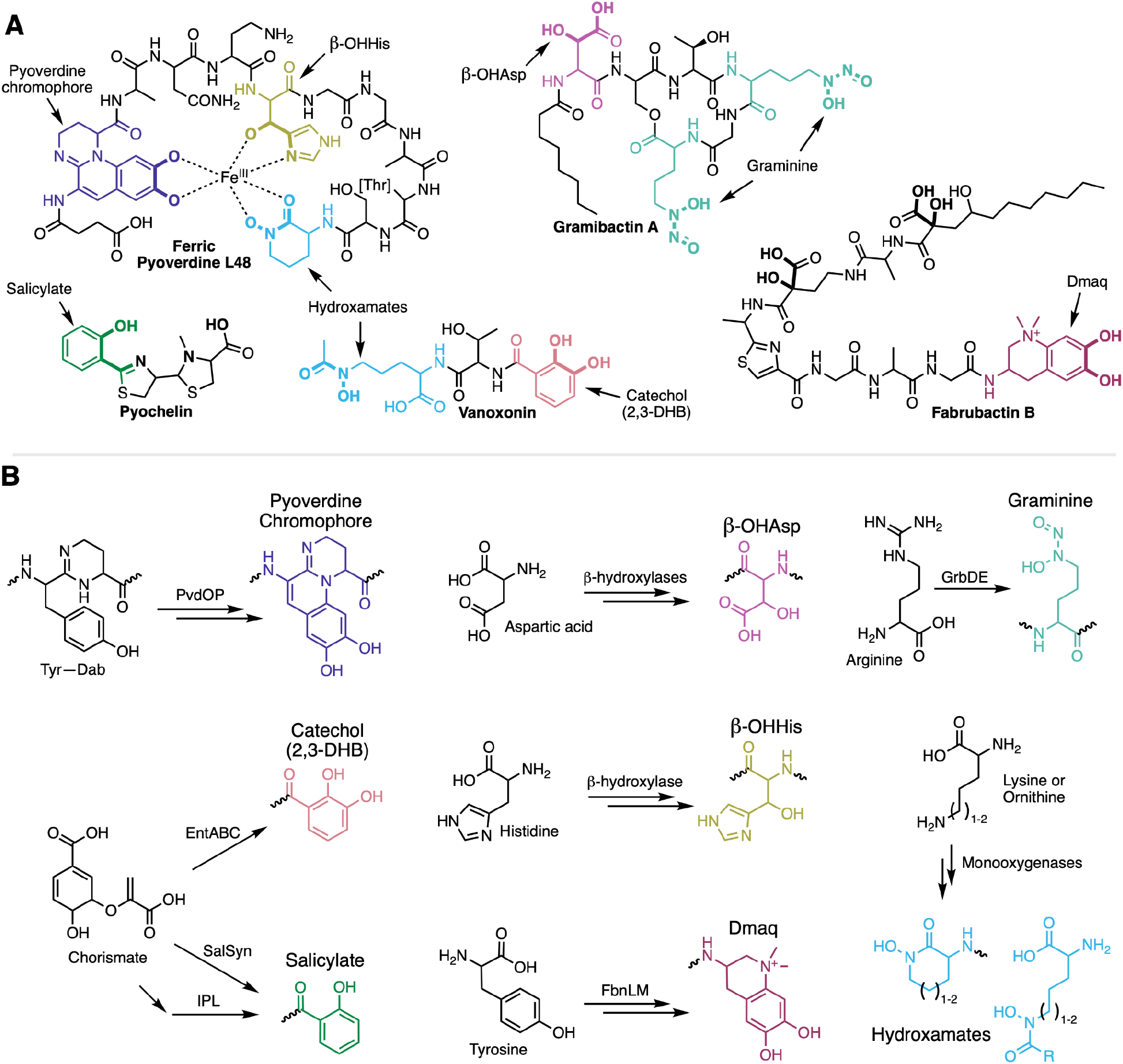
Chelating substructures found in bacterial NRP metallophores and their biosynthetic pathways. (A) Representative NRP metallophore structures. Nearly all known NRP metallophores contain one or more of the eight labeled chelating groups. Most chelating groups provide bidentate metal chelation, as shown for ferric pyoverdine L48. (B) Chelator biosynthesis pathways that form the basis for the new antiSMASH detection algorithm, as described in the text.

Mining genomes for metallophore BGCs has facilitated the discovery of chemically and biologically diverse metallophore systems; however, automated detection tools are still severely lacking.^9^ The peptidic backbones of NRP metallophores are produced by non-ribosomal peptide synthetases (NRPSs), large multi-domain enzymes that activate and condense amino acids and other substrates in an assembly-line manner.^11^ In the past two decades, a variety of bioinformatic tools have been developed to identify NRPS BGCs in a genome. One of the most popular is the secondary metabolite prediction platform antiSMASH, which uses a library of profile hidden Markov models (pHMMs) to identify (combinations of) enzyme-coding genes that are indicative of certain classes of specialized metabolite biosynthetic pathways.^12^ For example, antiSMASH identifies an NRPS BGC region by the minimum requirement of a gene containing at least one condensation and one adenylation domain. NRP metallophore BGCs are technically detected by this rule as well; however, NRPSs also produce many other families of compounds, and additional manual annotation is still required to identify NRP metallophore BGCs specifically. The current version of antiSMASH (6.2) has a “siderophore” rule; however, it only detects a single, unrelated family of enzymes, called NRPS-independent siderophore (NIS) synthetases.^13^ Thus, accurate prediction of BGCs encoding siderophores and other metallophores is currently limited to experts in natural product biosynthesis, and even experts cannot feasibly annotate hundreds or thousands of BGCs produced by high-throughput metagenomic or comparative genomic analyses. To date, no global analysis of NRP metallophores has been performed, and thus the prevalence, combinatorics, and taxonomic distribution of different chelating groups is unknown.

Here, we describe a method for the automated detection of NRP metallophore BGCs, using the presence of chelator biosynthesis genes within NRPS BGCs as key markers for predicting metallophore production. We have formalized and automated this strategy by developing a set of pHMMs and logical rules that detect biosynthetic pathways for eight different chelating groups found in NRP metallophores (Figure 1). Against a manually curated NRPS dataset, 78% of NRP metallophore BGCs were detected with a 3% false positive rate. The new detection rules were applied to 15,562 representative bacterial genomes, allowing us to take the first census of NRP metallophore production across bacteria. At least 25% of all NRPS clusters in representative genomes code for the production of metallophores. BGC similarity networking of 2,523 metallophore BGCs via BiG-SCAPE and analysis of combinatorial patterns of chelating groups revealed that significant biosynthetic diversity remains unexplored, with hundreds of NRP metallophore gene clusters unique to single strains in our dataset. By mapping NRP metallophore BGCs to the Genome Taxonomic Database (GTDB) phylogeny, we found that biosynthetic novelty can be found across many taxa, although metallophores from Cyanobacteria deserve greater attention. NRP metallophore detection is available in antiSMASH v7.0 through the web server (https://antismash.secondarymetabolites.org) and standalone command line tool.

## Results

### A chelator-based strategy for detection of NRP metallophore biosynthetic gene clusters

The specialized chelating moieties found in NRP metallophores are rarely found in other natural products, and thus we sought to automate metallophore BGC prediction by searching for genes encoding their biosynthesis. An extensive review of published NRP metallophore structures revealed that nearly all contain one or more of just eight chelator substructures (Figure 1A). Protein domains responsible for their biosyntheses have been reported (Figure 1B), and thus pHMMs could be constructed to allow detection of putative chelator biosynthesis genes. Generally, draft pHMMs were built from alignments of known and predicted NRP metallophore biosynthesis genes collected from literature (Figure S1). Initial bitscore cutoffs were determined by scanning MIBiG 2.0^14^ gene clusters, as well as 38 additional experimentally characterized NRP metallophore BGCs that we added to MIBiG 3.0 (Supplemental Table 1).^15^ Final cutoffs were determined by scanning 28,688 NRPS cluster regions from the antiSMASH database and manually inspecting hits near the initial cutoff to determine if they are likely true NRP metallophore BGCs based on the presence of genes encoding membrane transport, metal acquisition (ex: ferric reductases^16–18^), and multiple chelating groups. The pHMMs were iteratively refined where required by adding additional low-scoring putative true positives to the seed alignments until clear bitscore separations appeared between (putative) true and false hits (Figure S1). The final multiple sequence alignments, pHMMs, and cutoffs are provided in the Supplemental Dataset. A full description of each biosynthetic pathway detection strategy, including caveats and known limitations, is provided below. The profile HMMs found in antiSMASH are given in monospaced bold font.

### Detection of catechols and phenols

Like the aromatic amino acids, 2,3-dihydroxybenzoic acid (2,3-DHB) and salicylic acid are derived from chorismate (Figure 1B).^19–21^ The three-step synthesis of 2,3-DHB is catalyzed by an isochorismate synthase (**EntC**), an isochorismatase (EntB), and a 2,3-dihydro-2,3-dihydroxybenzoate dehydrogenase (**EntA**).^19^ The sole presence of **EntA** matches in a cluster was sufficient to detect 2,3-DHB-containing metallophore BGCs; however, only **EntC** could be used to eliminate the nearly-identical 3-hydroxyanthranilic acid pathway,^22^ and thus both **EntA** and **EntC** were required to accurately detect 2,3-DHB production. Salicylate biosynthesis was detected by the presence of either an isochorismate pyruvate-lyase (**IPL**)^20^ or a bifunctional salicylate synthase (**SalSyn**).^21^ Two condensation domain subtypes specific to catecholic and phenolic metallophores were also included as independent detection rules: VibH-like enzymes (**VibH**), which condense 2,3-DHB to diamines and polyamines,^23,24^ and tandem heterocyclization domains (**Cy_tandem**) are sometimes responsible for Ser, Thr, or Cys cyclization.^25^

### Detection of β-hydroxyamino acids

An analysis of siderophores containing β-hydroxy-aspartate (β-OHAsp) and β-hydroxy-histidine (β-OHHis) delineated three families of siderophore-specific Fe(II)/α-ketoglutarate-dependent enzymes responsible for β-hydroxylation of Asp (**TBH_Asp** and **IBH_Asp**) or His (**IBH_His**).^26^ More recently, β-OHAsp-containing siderophores named cyanochelins were isolated from a diverse group of cyanobacteria.^27^ The BGCs contain two additional classes of β-hydroxylases, tentatively named **CyanoBH_Asp1** and **CyanoBH_Asp2**. Because of the polyphyletic nature of these hydroxylases, well-performing pHMMs could not be made using the standard workflow used for other pathways (Figure S1); even after many iterations, non-metallophore NRPs were captured as false positives, such as the β-OHAsp-containing lipopeptide turnercyclamycin.^28^ Instead, pHMM construction was guided by a phylogenetic tree of β-hydroxylases from NRPS BGCs (Figure S2). Clades likely to be involved in metallophore biosyntheses were located using known genes from reported BGCs,^26,27^ and by the presence of nearby metallophore-specific transporter families (*vide infra*). Several unreported clades of amino acid β-hydroxylases were identified that may also be involved in metallophore biosynthesis based on their co-occurrence with transporter genes (Figure S2). However, these putative β-hydroxylases were not included in the current NRP metallophore rules due to a lack of experimental evidence for metallophore production. A negative constraint (in the form of a competing pHMM for **SBH_Asp**) was used to exclude the syringomycin family of BGCs, which contain a clade of β-hydroxylases that sit within the IBH_Asp clade.^26^

### Detection of hydroxamic acids

Hydroxamates are all produced by the hydroxylation and acylation of a primary amine. In peptidic metallophores, the amine is usually the sidechain of ornithine (Orn) or Lys, which is first hydroxylated by a flavin-dependent monooxygenase (**Orn_monoox** or **Lys_monoox**, respectively). Vicibactin is an unusual hydroxamate siderophore^29^ that could not be accurately captured by either **Orn_monoox** or **Lys_monoox**. The Orn hydroxylase VbsO is more similar to those found in NIS pathways, and a VbsO pHMM was non-specific. Instead, the acyl-hydroxyornithine epimerase^29^ **VbsL** is used to detect vicibactin. In two non-metallophore pathways,^30,31^ hydroxyornithine or hydroxylysine are the substrates for N-N bond-forming enzymes. No clear bitscore separation could be achieved to eliminate these false positives, so two negative constraints were added: the presence of **KtzT** associated with biosynthesis of piperazates,^30^ and **MetRS-like**, associated with several hydrazines.^31^ Unfortunately, false positives may still arise if the ornithine monooxygenase and the negative constraint genes are (distantly) located within the same BGC region (>30 kbp), such as in the himastatin locus (MIBiG BGC0001117).

### Detection of other chelating groups

Although the biosynthesis of the chelating amino acid graminine has not been fully elucidated, gene knockouts and stable isotope studies have revealed two enzymes, **GrbD** and **GrbE**, responsible for diazeniumdiolate formation from arginine.^32,33^ The quinoline chelator Dmaq was first identified in anachelins,^34^ and the biosynthesis was recently established for fabrubactins (Figure 1).^35^ Synthesis is initialized by **FbnL** and **FbnM**, which form a two-protein heme peroxidase that oxidizes L-tyrosine to L-DOPA.^35^ GrbDE and FbnLM were previously used as a handle for genome mining,^32,35^ and our pHMMs gave similar results. The diverse pyoverdine family of siderophores is defined by a fluorescent chelating chromophore (Figure 1). Chromophore maturation is driven by the tyrosinase **PvdP** and the oxidoreductase **PvdO**.^36,37^ The two genes are sometimes in separate loci, and including both as independent pHMMs increased the number of pyoverdine BGCs detected. These pHMMs also capture the biosynthesis pathway for azotobactin, which contains a slightly different chromophore (Figure 3).^38^

### Currently undetectable chelating groups

Biosynthetic pathways for a few NRP metallophore chelating groups could not be detected using the current strategy. The antiSMASH detection rules are generally limited to identifying protein subfamilies unique to metallophore biosyntheses and cannot detect chelators synthesized by the core NRPS and/or polyketide synthase (PKS) machinery without also retrieving many false positives. For example, NRPS-derived thiazol(id)ine and oxazol(id)ine heterocycles (see pyochelin, Figure 1A) are found in many disparate classes of NRPs in addition to metallophores. Most heterocycle-containing metallophore BGCs are still detected, as they contain a detectable salicylate or hydroxamate pathway, or are captured by the metallophore-specific **Cy_tandem** pHMM. However, the micacocidin family of metallophores is currently undetectable by the current algorithm, as their 5-alkylsalicylate chelating moiety is not derived from chorismate, but rather produced by an iterative PKS.^39^ A future antiSMASH metallophore module may be able to detect (combinations of) subclades of multiple NRPS and PKS domains to predict the biosynthesis of additional metallophore substructures.

Additionally, one rare pathway was not included: fabrubactins contain two α-hydroxycarboxylate chelating moieties not reported in any other metallophore (Figure 1A, bolded atoms).^35^ Both substructures are proposed to be synthesized by the flavin-dependent monooxygenase FbnE. A profile HMM was built for FbnE (Supplemental dataset) but was not included in antiSMASH, as all BGCs with **FbnE** hits were either captured independently by **FbnL** and **FbnM** (Dmaq biosynthesis), or appeared to be false positives based on a lack of other metallophore-related genes.

### Validating antiSMASH NRP metallophore detection against manually curated BGCs

In order to assess the performance of the NRP metallophore BGC detection strategy developed herein, we manually predicted metallophore production among a large set of BGCs. A total of 758 NRPS BGC regions from 330 genera were annotated with default antiSMASH v6.1 and inspected for markers of metallophore production, including genes encoding transporters, iron reductases, chelator biosynthesis, and known metallophore NRPS domain motifs. We thus manually classified 176 BGC regions (23%) as metallophore BGCs (Supplemental Table 2). The new antiSMASH detection rules were applied to the same BGC regions, resulting in 145 putative metallophore BGC regions (F1 = 0.86; Table 1 and Supplemental Table 2). Nine metallophore BGC regions were only detected by antiSMASH. Upon reinvestigation, four were determined to be genuine metallophore BGC regions missed during manual analysis, leaving only five putative false positives in which seemingly unrelated genes matched the pHMMs (97% precision). Conversely, a total of 40 metallophore BGC regions could only be detected manually (78% recall). Two of the false negatives encoded the 5-alkylsalicylate PKS that is currently undetectable, as described above. In the majority of false negatives, NRP metallophore BGCs were missed because chelator biosynthesis genes, on which the detection strategy is based, were not present in the cluster. In 21 cases, genes encoding catechol, salicylate, or hydroxamate biosynthesis were located elsewhere in the genome. In ten cases, BGCs had NRPS domains for the incorporation of catechol, salicylate, or hydroxamate residues, but chelator biosynthesis pathways were not found anywhere in the genome; these clusters may be non-functional fragments, rely on exogenous precursors (as seen in equibactin biosynthesis^40^), or have evolved to use novel chelator biosyntheses. Finally, seven manually assigned NRP metallophore BGC regions had no genes corresponding to known chelator pathways; if correctly annotated, they may represent novel structural classes.

**Table 1.** Performance metrics for NRP metallophore detection. The chelator-based rules newly implemented in antiSMASH, the transporter-based method of Crits-Christoph *et al*.,^41^ and a combined either/or ensemble were each tested on a set of 758 NRPS BGC regions that were manually annotated to find 180 metallophore BGCs. Full results are given in Supplemental Table 2. ^†^F1 score is equal to 2×(Precision×Recall)/(Precision+Recall).

|  | Precision | Recall | F1<sup>†</sup> |
| --- | --- | --- | --- |
| AntiSMASH rules | 0.97 | 0.78 | 0.86 |
| Transporter genes | 0.93 | 0.56 | 0.69 |
| Either/or ensemble | 0.92 | 0.88 | 0.90 |

One particularly promising novel BGC was found in the genome of *Sporomusa termitida* DSM 4440, an anaerobic acetogen (Figure S3). The BGC was manually annotated as a putative metallophore due to the presence of a TonB-dependent outer membrane receptor, as well as NRPS domains, methyltransferases, and oxidoreductases homologous to those involved in the biosynthesis of pyochelin and other thiazol(id)ine metallophores. A salicylate synthase gene, present in similar clusters, appears to have been replaced by a partial menaquinone pathway (*menFDEB*). We thus predicted that the salicylate moiety would be replaced by 1,4-dihydroxy-2-naphthoic acid (Figure S3). The Natural Product Atlas contained one family of bacterial compounds with that substructure, karamomycins, which were isolated from an unsequenced strain of *Nonomuraea endophytica* (Figure S3).^42^ Karamomycins also contain thiazol(id)ines, as predicted for the *S. termitida* DSM 4440 BGC product. Hence, karamomycins are likely produced by a homologous BGC, and both compounds may be involved in trace metal binding and transport.

### AntiSMASH outperforms transporter-based detection, although both techniques are complementary

Crits-Christoph et al. found that the presence of transporters could be used to predict siderophore BGCs among other NRPS clusters.^41^ Specifically, the Pfam families for TonB-dependent receptors, FecCD domains, and periplasmic binding proteins (PF00593, PF01032, and PF01497, respectively) were determined to be highly siderophore-specific, and the authors used the presence of two of the three domains to predict a “siderophore-like” BGC region (metallophores that transport other metals were also coded as siderophores in their dataset). We used a modified version of antiSMASH to detect the three transporter families among the 758 NRPS BGC regions that we manually annotated for metallophore production (Table 1 and Supplemental Table 2). In total, the transporter-based method detected 108 metallophore clusters (F1 = 0.69), including eight putative false positives (93% precision), and had 80 false negatives (56% recall). One false positive was noted in the manual annotation as a likely “cheater”: while several *Bordetella* genomes encode the synthesis of a putative graminine-containing metallophore, *B. petrii* DSM 12804 has retained only the transporter genes alongside a small fragment of the NRPS. In the seven other false positives, BGC regions appeared to coincidentally contain transporter genes in their periphery, as they were not conserved in homologous NRPS clusters. In one case, the triggering genes were part of a vitamin B12 import and biosynthesis locus. Combining the two methods in an either/or ensemble approach slightly improved overall performance *versus* the antiSMASH rules alone, achieving 92% precision, 88% recall, and an F1 score of 0.90 (Table 1).

### Charting NRP metallophore biosynthesis across bacteria

The implementation of NRP metallophore BGC detection into antiSMASH allowed us to take the first bacterial census of NRP metallophore biosynthesis. The finalized detection rules were applied to 15,562 representative bacterial genomes from NCBI RefSeq (25 June 2022). In total, 3,264 NRP metallophore BGC regions were detected (Table 2 and Supplemental Table 3), including 54 Type II (non- or semi-modular^43^) NRPS regions that would otherwise not be detected by antiSMASH, such as BGCs for acinetobactin and brucebactin.^44,45^ NRP metallophores comprised 16% of all NRPS BGC regions in the genomes. Among complete regions (not located on a contig edge), 21% of NRPS BGC regions were classified as NRP metallophores, compared to just 8.6% of incomplete NRPS regions. This is consistent with previous reports that low-quality, fragmented genomes result in low-quality BGC annotations in antiSMASH.^46^ The transporter-based approach predicted siderophore activity for 15% of complete NRPS regions, including 463 BGC regions without detectable chelator genes; when the two methods are combined, over 25% of NRPS BGCs are predicted to produce NRP metallophores (Table 2). Only complete NRP metallophore BGC regions detected by antiSMASH were used for downstream analyses.

**Table 2.**
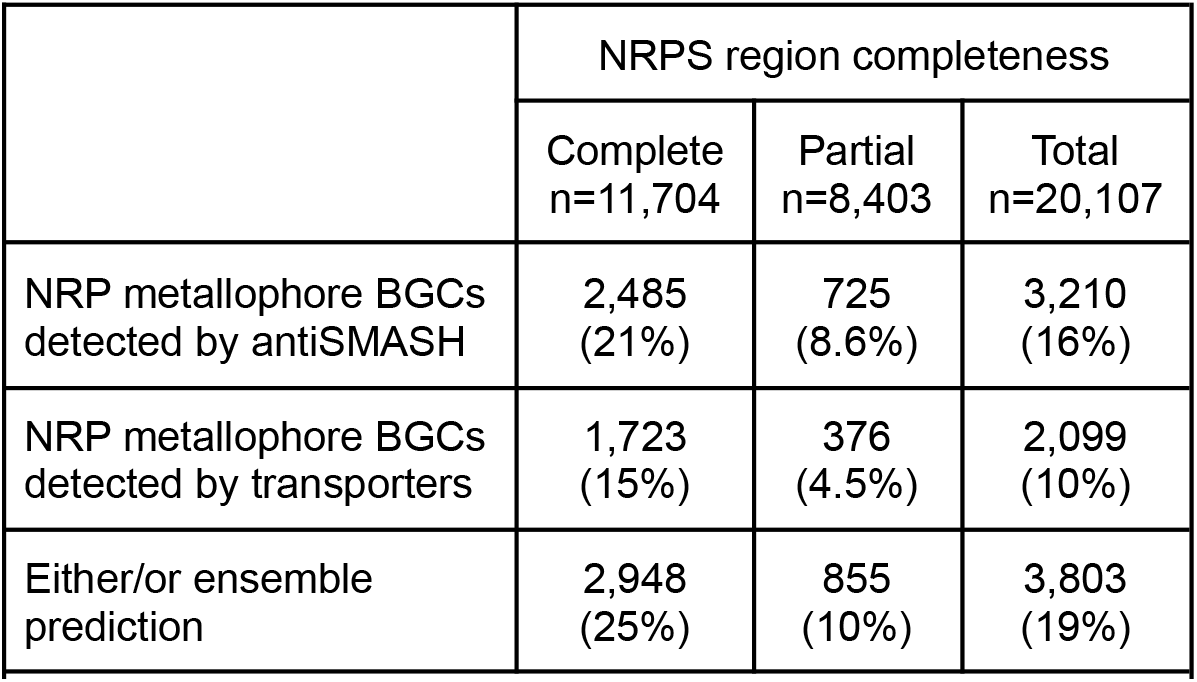
Summary of NRP metallophore BGC regions detected in representative bacterial genomes. The chelator-based rules newly implemented in antiSMASH, the transporter-based method of Crits-Christoph et al.,^41^ and a combined either/or ensemble were applied to 15,562 NCBI RefSeq representative bacterial genomes. The full results are given in Supplemental Table 3. A region is “complete” if it is not on a contig edge, as determined by antiSMASH. An additional 54 NRP metallophore BGC regions (38 complete) were detected that do not appear in this table because they did not meet the antiSMASH requirements for an NRPS BGC region (a single coding sequence with at least one condensation domain and adenylation domain).

### Frequency and hybridization of NRP metallophore chelating groups

Complete NRP metallophore regions from the representative genomes were categorized by the type(s) of chelator biosynthesis genes detected within (Figure 2). Hydroxamates and catechols were the most common pathways, present in 44% and 36% of BGC regions, respectively. In contrast, β-OHHis, graminine, and Dmaq biosyntheses were rare in representative genomes, each present in less than 2% of detected regions. About 20% of regions contained genes for at least two pathways and putatively produce a hybrid metallophore. Only 42 BGC regions (1.7%) contained three different chelating groups: each encoded genes for the pyoverdine chromophore, a hydroxamate, and either β-OHAsp or β-OHHis. The proportion of hybrid metallophores is likely higher than estimated here. As described above, some chelating moieties could not be captured by the pHMM-based rules. For example, many salicylate metallophores contain additional oxazolidine or thiazolidine metal-binding moieties (ex: pyochelin, Figure 1A) and all 15 BGC regions that are predicted to only encode the Dmaq pathway also have homologs of the α-hydroxycarboxylate synthase gene *fbnE*. Furthermore, metallophore biosynthesis may require genes from multiple BGCs. Pyoverdine genes may be located in up to five different loci,^47^ and all 56 regions with only the pyoverdine chromophore pathway are expected to produce hybrid siderophores. Two chelator combinations found in our dataset have not previously been reported in NRP metallophore BGCs: hydroxamate / β-OHHis hybrid BGCs were found in three *Pseudomonas* species, and a single putative hydroxamate / graminine hybrid was found in *Tistrella mobilis*. Excluding pyoverdines,^48,49^ exochelin MN is the only reported hydroxamate / β-OHHis hybrid metallophore, produced by an unsequenced *Mycobacterium neoaurum* strain.^50^ The predicted structures of the novel *Pseudomonas* hydroxamate / β-OHHis hybrids are given in Figure S4. All other chelator combinations in our dataset have been reported in characterized siderophores, as exemplified by the structures in Figure 3.

**Figure 2.**
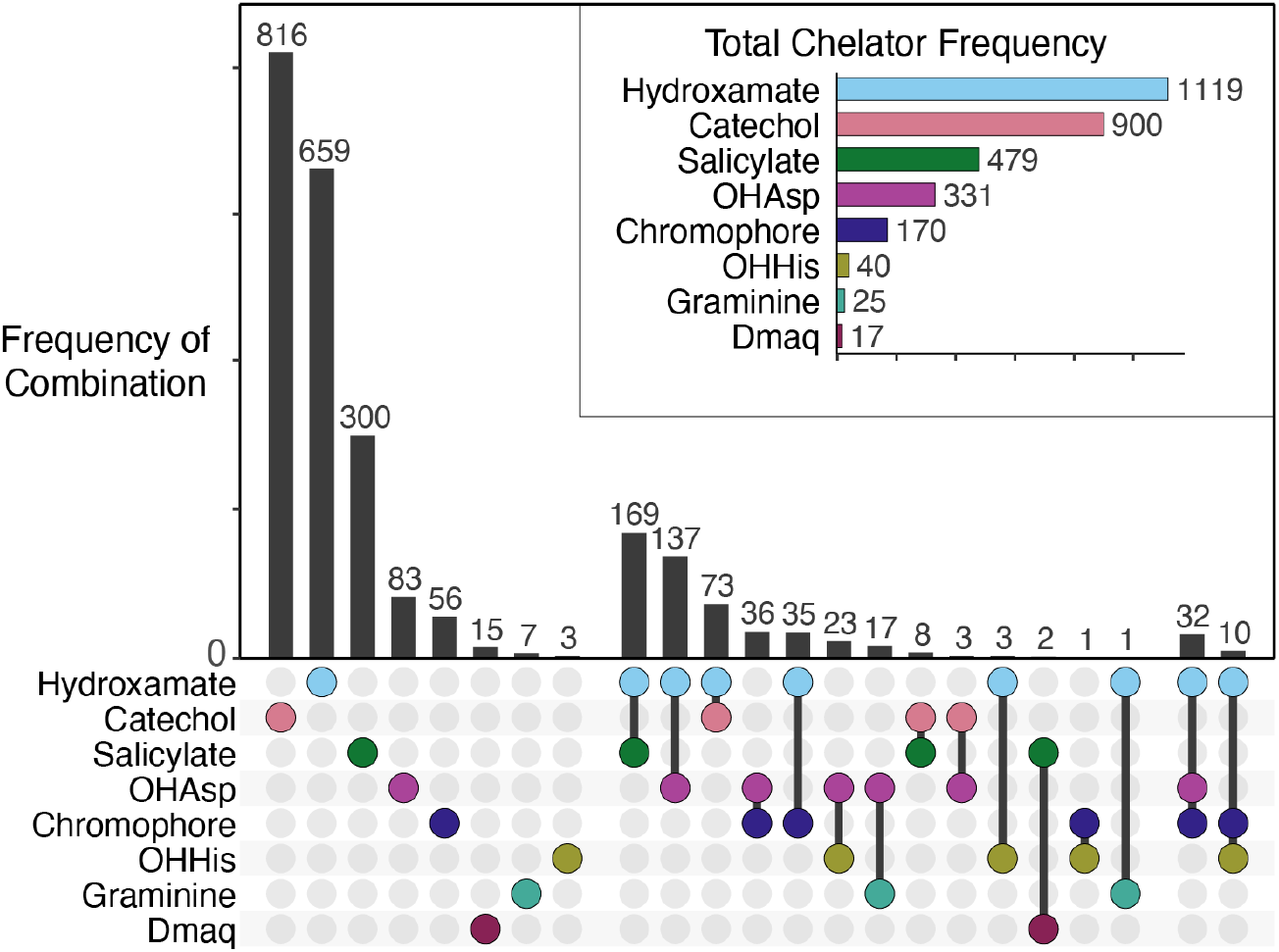
An upset plot of chelator frequency among 2,489 complete NRP metallophore BGC regions from RefSeq representative genomes. An additional 38 BGC regions were detected by metallophore-specific NRPS domains (VibH-like or tandem Cy) rather than chelator biosynthesis, and may produce catechol and/or salicylate metallophores using biosyntheses encoded elsewhere in the genome.

**Figure 3.**
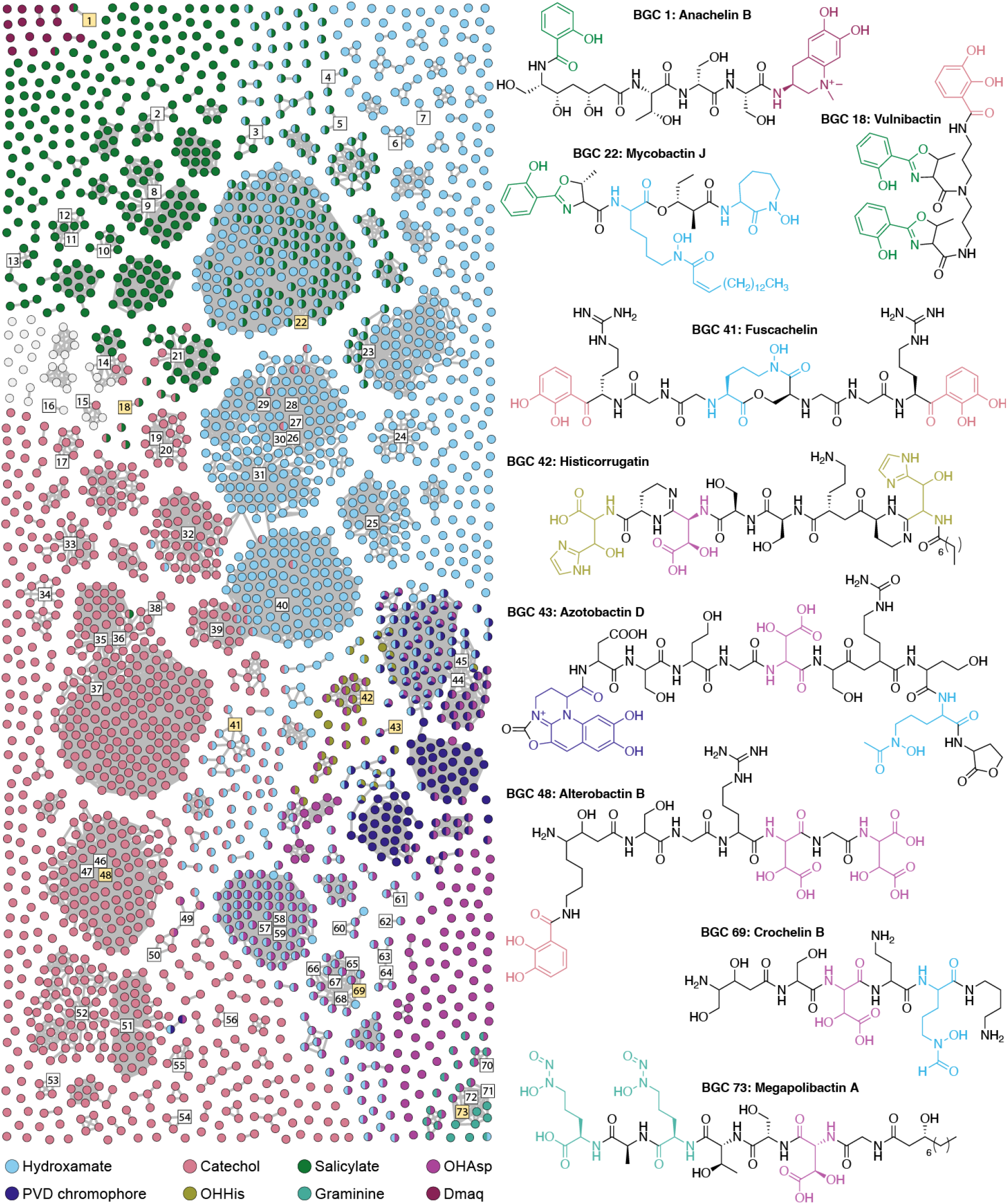
BiG-SCAPE similarity network of complete NRP metallophore BGC regions from RefSeq representative genomes. Numbered square nodes indicate published BGCs, as given in Supplemental Table 1. Select hybrid metallophore BGC nodes are highlighted yellow, and their corresponding structures are shown. Nodes are colored by the type(s) of chelator biosynthesis detected therein. BGC regions colored light gray contain only metallophore-specific NRPS domains (VibH-like or tandem Cy) and may produce catechol and/or salicylate metallophores using biosyntheses encoded elsewhere in the genome. The network was constructed in BiG-SCAPE v1.1.2 using 2,596 BGC regions as input, including 78 reference BGCs, and a distance cutoff of 0.5.

### The most widespread NRP metallophore families have likely been found, yet significant diversity remains unexplored

Different species of bacteria can contain highly similar metallophore BGCs. To gauge the biosynthetic diversity of the putative NRP metallophores (and thereby the structural diversity), the complete BGC regions were organized into a sequence similarity network using BiG-SCAPE, which clusters BGCs based on their shared gene content and identity. An additional 75 reported NRP metallophore BGCs were included as reference nodes (Supplemental Table 1), and a distance cutoff of 0.5 was chosen to allow highly similar reference BGCs to form connected components (gene cluster families; GCFs) in the network. The final network, colored and organized by chelator type, is presented in Figure 3. The majority of BGC regions (57%) clustered with the reference BGCs in just 45 GCFs, suggesting that many of the most widespread NRP metallophore families already have characterized representatives (Figure S5). However, 1093 BGC regions did not cluster with any reference BGC, forming 619 separate GCFs in the network (93% of all GCFs). Some of these may encode orphan metallophores previously isolated from unsequenced strains, or be similar to known BGCs that were not included in our non-exhaustive literature search. Nevertheless, significant NRP metallophore structural diversity remains undiscovered, particularly among the 484 BGC regions distinct enough to form isolated nodes in the network.

### NRP metallophores are rarely produced by anaerobic bacteria

Although the majority of known NRP metallophores are classified as siderophores, metallophores can transport many other metals, as well as play roles in oxidative stress response, cell signaling, and antibiotic activity.^51^ The presence of metal chelator biosynthesis genes in an NRPS BGC is therefore not sufficient to assign a specific biological role to the molecule(s) produced. Nevertheless, we expect that most detectable NRP metallophore BGCs produce siderophores (possibly in addition to other roles), given literature precedent and the high Fe(III) affinity of bidentate ligands with anionic oxygens.^1,2^ Siderophores are only rarely produced under anaerobic conditions due to the relatively high solubility of reduced Fe(II),^1,52,53^ and thus we hypothesized that NRP metallophore BGCs would be most prevalent in aerobic strains. Using oxygen tolerance data from BacDive^54^, 863 strains with complete RefSeq representative genomes were categorized as aerobes, anaerobes, or “other” (including facultative anaerobes and microaerophiles, as described in Methods). Only 2.2% of anaerobes carried detectable NRP metallophore BGCs, compared to 21% of aerobes (Fisher’s exact test, *p*=1e-13). Considering only strains with at least one NRPS BGC, 13% of anaerobes encoded an NRP metallophore, compared to 50% of aerobes (*p*=8e-6). Because of phylogenetic structure, related species do not totally satisfy the independence criterion of Fisher’s exact test; however, the significance held when sampling one strain per genus (*p*=2e-6) and one strain per genus with at least one NRPS BGC (*p*=0.02). Of the six metallophore BGC regions from anaerobic strains, five putatively encode the synthesis of yersiniabactin-like compounds. Uropathogenic *E. coli* (facultative anaerobe) uses yersiniabactin, a broad-spectrum metallophore, to import nickel and prevent host-induced copper toxicity;^55,56^ the BGCs found herein may serve similar non-ferric roles. The sixth strain, *Blastochloris viridis*, was coded as an anaerobe by BacDive; however, it is capable of microaerophilic growth.^57^ The putative turnerbactin-like catechol siderophore may be involved in protecting *B. viridis* from oxidative stress^58^ or be involved in solubilizing Fe(III)-oxides.^53^

### NRP metallophore biosynthesis is unevenly distributed across bacterial taxa

In order to highlight understudied metallophore producers, taxonomic distribution of detectable NRP metallophore biosynthesis was gauged using the bacterial reference tree produced by the Genome Taxonomy Database^59^ (Figure 4). The tree taxa were pruned to include only the 2,195 genomes that were also RefSeq representatives and complete genomes, and then antiSMASH results were mapped onto the tree using the Interactive Tree Of Life (iTOL).^60^ Predicted NRP metallophore BGC regions were found in 15% of mapped genomes (328 / 2,195), nearly all of which belong to just five phyla: at least one BGC was found in 60% of Myxobacteria, 41% of Cyanobacteria, 35% of Actinobacteria, 18% of Proteobacteria, and just 3.3% of Firmicutes (Figure 4). The phyla Gemmatimonadota and Nitrospirota each contained one mapped strain with a predicted NRP metallophore BGC: *Gemmatimonas aurantiaca* putatively produces enterobactin, and *Nitrospira japonica* putatively produces a novel β-OHAsp-containing siderophore (Figure 4). The prevalence of NRP metallophore BGCs also varied significantly across proteobacterial and actinobacterial orders (chi square test, *p*<1e-15 and *p*=1e-12, respectively). For example, only 5.4% of Actinomycetales genomes contained an NRP metallophore BGC, compared to 59% of Mycobacteriales genomes and 52% of Streptomycetales genomes (Fisher’s exact test, *p*<1e-15 and *p*=4e-10, respectively). The uneven distribution of NRP metallophores can only partially be attributed to the uneven distribution of NRPS systems themselves (Figure S6). Among genomes with at least one modular NRPS BGC detected by antiSMASH (n=657), 49% contained NRP metallophores on average, with significant variation across phyla (Fisher’s exact test, Bonferroni correction): predicted NRP metallophore clusters were not found in Bacteroidota and Planctomycetota (*p*=1e-14, *p*=1e-3), rare in Firmicutes (in 19% of NRPS-containing strains, *p*=6e-7), and abundant in Actinobacteriota (in 68% of NRPS-containing strains, *p*=8e-10). Predicted NRP metallophore BGCs were found in the majority of Myxococcota and Cyanobacteria strains with at least one NRPS BGC (75% and 65%, respectively); however, few strains had been retained after filtering, and the overabundance was not statistically significant. Dedicated analyses of Myxococcota and Cyanobacteria with larger sample sizes will be required to thoroughly explore their NRP metallophore prevalence and diversity.

**Figure 4.**
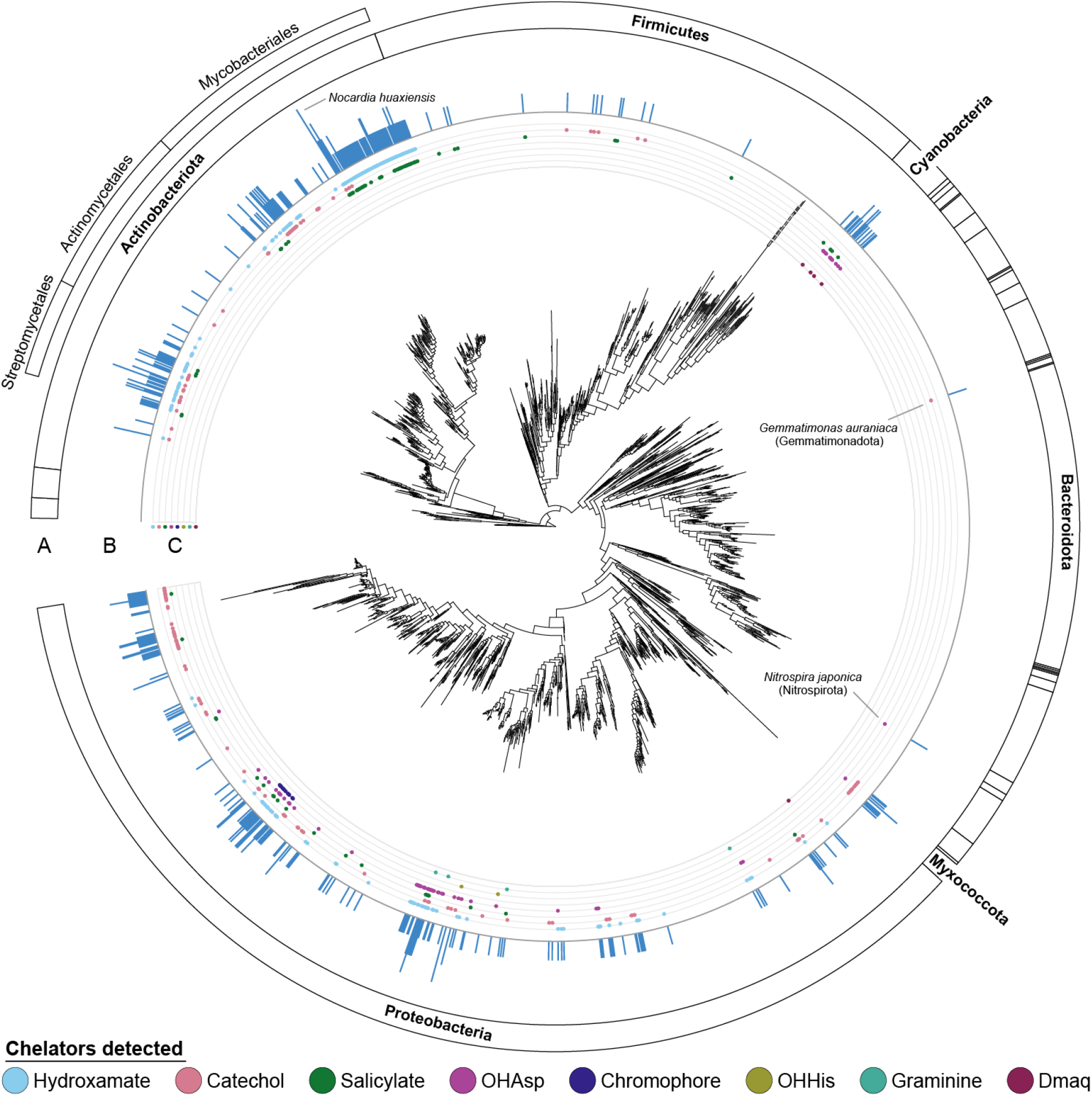
NRP metallophore biosynthesis, mapped to the Genome Taxonomy Database (GTDB) phylogenetic tree. Ring A: Phylal divisions. Key phyla are labeled, as well as Actinobacteriota orders mentioned in the text. Ring B: The number of NRP metallophore BGCs detected in each genome. *Nocardia huaxiensis* contained four putative BGCs, the most detected in any genome. Ring C: Chelator biosynthetic pathways detected in each genome. Center: The GTDB species tree (version r207), pruned to complete RefSeq representative genomes. Annotations were mapped to the phylogenetic tree using iTOL v6.6.

BiG-SCAPE networking had revealed that putatively novel BGCs (dissimilar from reference BGCs) comprised 43% of NRP metallophore BGCs in representative genomes (Figure 3 and Figure S5). To identify taxa harboring untapped NRP metallophore biosynthetic diversity, we mapped GCF novelty from BiG-SCAPE onto the GTDB phylogenetic tree in Figure 4 using iTOL (Figure S7). The phyla Actinobacteriota, Proteobacteria, Firmicutes, and Myxococcota showed no significant difference in their proportions of novel NRP metallophore BGCs (40-50%), while 94% of Cyanobacterial NRP metallophore BGCs were novel (17 of 18, *p*=2e-4). Myxococcia was the only bacterial class that had NRP metallophore BGCs but no novel GCFs; all clusters belonged to the myxochelin GCF (Figure 3, nodes 19 and 20). In contrast, the sister class Polyangia contained only novel NRP metallophore BGCs, which comprised five singleton GCFs in the BiG-SCAPE network. Four of the five BGCs from Polyangia were visually similar in gene content to the myxochelin BGC, and likely produce (potentially novel) myxochelin congeners. Although most taxa with NRP metallophore BGCs contained novel GCFs, even for well-studied genera, manual inspection will be required to determine if the BGCs truly correspond to new structural diversity.

Of the eight detectable NRP metallophore chelating groups (Figure 1), only two are synthesized by the same protein family (β-OHAsp and β-OHHis), and therefore, metal-chelating moieties must have evolved multiple times. Accordingly, each chelating pathway had a distinct taxonomic distribution among mapped genomes (Figure 4, ring C). Hydroxamate, catechol, salicylate, β-OHAsp, and Dmaq chelating pathways were detected in multiple phyla; while graminine, β-OHHis, and the pyoverdine chromophore were limited to select Proteobacteria. Furthermore, distinct distributions were also observed within hydroxamate, β-OHAsp, and salicylate biosynthesis, each of which is produced by multiple clades within the same enzyme family (Figure S8). No chelator pathway was consistently present across its taxonomic range. Certain lineages may have lost pathways from their genomes in favor of xeno-metallophore usage or replacement by another metallophore pathway, and other lineages may have acquired pathways from other organisms by horizontal gene transfer. Both of these events have been reported among siderophores.^61,62^ Comprehensive phylogenetic analyses of each chelating group are underway, which will help elucidate their evolutionary histories.

## Discussion

Trace metal starvation shapes interactions within microbial communities and between bacteria and the host; therefore, natural and synthetic microbiomes cannot be understood without knowing the metallophore biosynthetic potential of the community. High-throughput biotechnological applications will benefit from *in silico* metallophore prediction due to the prohibitively high cost of isolation and characterization. To date, distinguishing peptidic metallophore BGCs from other NRPS BGCs has been largely limited to manual expert analyses, leading to blind spots in our understanding of microbes and their communities. We have now automated bacterial NRP metallophore prediction by extending the secondary metabolite prediction platform antiSMASH to detect genes involved in the biosynthesis of metal chelating moieties (Figure 1). The new detection strategy was applied to 15,562 representative bacterial genomes, and 3,264 putative NRP metallophore BGC regions were found in total. In our initial investigations into this rich dataset, we conservatively estimate that one fourth of NRPS BGCs encode metallophore production, predict new combinations of chelating groups, statistically support the relationship between aerobic bacteria and siderophore production, and identify Cyanobacteria as prime targets for the isolation of novel metallophores. The NRP metallophore detection algorithm is publicly available in the antiSMASH v7.0 web server and command line tool (https://antismash.secondarymetabolites.org).

The presence of genes encoding catechols, hydroxamates, and other chelating groups is one of the most frequently used markers of a metallophore BGC.^9^ We have formalized and automated the identification of eight chelator pathways, allowing us to detect 78% of NRP metallophore BGCs with a 3% false positive rate against a manually annotated set of NRPS clusters. Biosynthetic genes are detected with custom pHMMs and significance score cutoffs calibrated for accurate metallophore discovery, diminishing the ambiguity of interpreting gene annotations, protein families, and BLAST results. We acknowledge that human biases may have influenced which clusters were coded as putative metallophores during both algorithm development and testing; however, expert manual curation remains the most reliable method for NRP metallophore BGC detection. During testing, antiSMASH correctly identified four NRP metallophore BGCs that were initially missed by manual curation, suggesting that future algorithms, incorporating additional forms of evidence, will be able to outperform researchers. Unfortunately, 22% of manually identifiable metallophore BGCs could not be automatically distinguished from other NRPS clusters. The rules developed herein rely on the presence of one or more known chelator biosynthesis genes colocalized with the NRPS genes. In some cases, biosynthetic genes are located elsewhere in the genome, for example in a second NRP metallophore cluster. Advances in the machine-learning prediction of adenylation domain specificity should aid in identifying NRPS modules that incorporate chelating moieties. Additionally, some substructures capable of metal chelation are produced by the core NRPS/PKS machinery rather than unique accessory enzymes, including 5-alkylsalicylates and oxazolidines. The current algorithm can detect two metallophore-specific NRPS condensation domain subtypes (VibH-like and Cy_tandem), and phylogenetic analyses may reveal more metallophore-specific clades of NRPS or PKS domains.

Recently, Crits-Christoph et al. demonstrated the use of transporter families to predict that a BGC encodes siderophore (or metallophore) biosynthesis.^41^ Among our test dataset, the biosynthesis-based antiSMASH rules outperformed the transporter-based approach (F1 = 0.86 *versus* F1 = 0.69). However, some putative metallophore BGCs were only found using the transporter-based approach, and a combined either/or ensemble approach slightly outperformed the antiSMASH rules alone (F1 = 0.90). Biosynthetic- and transporter-based techniques are thus complementary, and future work will incorporate transporter genes into antiSMASH metallophore prediction. We note that the reported transporter-based approach uses just three pHMMs from Pfam, while our biosynthetic detection requires many custom pHMMs. An extended set of metallophore-specific transporter pHMMs designed according to the same principles as those followed for the biosynthesis-related pHMMs could significantly improve detection by reducing false positives and capturing other families of transporters. The NRP metallophore BGCs discovered in this study could serve as a dataset for developing a more comprehensive model for metallophore transporter detection.

Siderophore and other metallophore structures are often broadly categorized by their chelating group, as each chelator has unique chemical properties (ex: hydrophobicity, base hardness, pKa, photoreactivity) that directly influence their metal binding and biological role. We likewise categorized 2,523 NRP metallophore BGCs according to the chelator biosynthesis contained within (Figure 2). Chelator pathway abundance was consistent with reported NRP metallophore structural diversity: hydroxamate and catechol biosyntheses were common in representative genomes, while β-OHHis, graminine, and Dmaq were rare. Hybrid metallophore BGCs, with more than one detectable chelator biosynthesis pathway, were found to make up 20% of NRP metallophores. This is likely an underestimate, as certain chelating groups and multi-locus BGCs currently cannot be detected. The diverse enzyme families responsible for the biosynthesis of NRP metallophore chelating groups (Figure 1B) evince that metal chelation has evolved multiple times, and we expect that more NRPS chelator substructures remain undiscovered. In fact, during manuscript preparation, the novel chelator 5-aminosalicylate was reported in the structure of the *Pseudonocardia* NRP siderophore pseudonochelin,^63^ and we found several unexplored clades of Fe(II)/α-ketoglutarate-dependent amino acid β-hydroxylases that are likely involved in metallophore biosyntheses (Figure S2). Additionally, we have likely identified a new biosynthetic pathway in the genome of *Sporomusa termitida* DSM 4440, which encodes a partial menaquinone pathway in place of a salicylate synthase to putatively produce a novel karamomycin-like metallophore (Figure S3).^42^ The modular nature of the pHMM-based detection rules will allow for new chelating groups to be added as their biosyntheses are experimentally characterized.

Metallophore BGC regions from representative genomes were compared to reference BGCs and organized into gene cluster families (GCFs) with BiG-SCAPE (Figure 3). Only 7% of GCFs contained a characterized reference genome, yet these GCFs include the majority of detectable NRP metallophore BGCs. We found 1093 metallophore BGC regions that were dissimilar from any reference BGC, and almost 500 distinct BGC regions were found in only a single strain. Although significant biosynthetic diversity remains undiscovered, cluster de-replication will become increasingly important to avoid re-isolating known compounds. Unfortunately, the BiG-SCAPE analysis was constrained by an incomplete list of known BGCs. For this work, we added 37 NRP metallophore BGCs to MIBiG v3.0, doubling the database’s coverage (Supplemental Table 1), and yet many published BGCs have yet to be added to the database. Efforts are currently underway to computationally deorphanize reported metallophore structures and connect them to BGCs detected herein.

Detectable NRP metallophore BGCs were mapped onto the bacterial species phylogeny from GTDB to highlight taxa for future study. Consistent with reported NRP metallophore characterization, BGCs were largely constrained to the phyla Actinobacteria, Proteobacteria, Cyanobacteria, Firmicutes, and Myxobacteria. We found that Cyanobacteria genomes contain novel, diverse BGCs and deserve dedicated campaigns to characterize their metallophore diversity, although putatively novel siderophores were found across many taxa. Each NRP siderophore chelator biosynthesis pathway was found to have a unique taxonomic distribution, consistent with a complicated evolutionary history where each chelator evolved independently and has since been horizontally transferred, lost to cheating, and combined into hybrid siderophores; thus producing the NRP siderophore diversity seen today. Comparing gene and species phylogenies may help age determination of each siderophore chelating group, and reveal if any of the chelators found in multiple phyla are more ancient than phyla themselves.

In this work, we have taken a major step towards accurate, automated NRP metallophore BGC detection. The new strategy affords a clear improvement over manual curation, and has already allowed for the high-throughput identification of thousands of likely NRP metallophore BGC regions. A future antiSMASH module might predict metallophore activity more accurately with a machine learning algorithm that considers multiple forms of genomic evidence, including the presence of transporter genes, NRPS domain architecture and sequence, metal-responsive regulatory elements, and other markers of metallophore biosynthesis that are still limited to manual inspection.^9^ In particular, regulatory elements will likely be required to accurately distinguish siderophores, zincophores, and other classes of metallophores. Implementation of NRP metallophore BGC detection into antiSMASH will enable scientists of diverse expertises to identify and quantify NRP metallophore biosynthetic pathways in their bacterial genomes of interest and promote large-scale investigations into the chemistry and biology of metallophores.

## Methods

For all software, default parameters were used unless otherwise specified. All python, R, and bash scripts used in this paper, as well as underlying data, is available in the Supplemental Dataset, published to Zenodo: 10.5281/zenodo.7439477.

### Profile hidden Markov model construction

Profile hidden Markov models (pHMMs) were built from biosynthetic genes in known metallophore pathways, supplemented with putative BGC genes where required (Figure S1 and Supplemental Dataset, 1_development/). Amino acid sequences were aligned with MUSCLE (v3.8)^64^ and pHMMs were constructed using hmmbuild (HMMER3).^65^ pHMMs were tested against the MIBiG database (v2.0)^14^ and an additional 37 NRP siderophore BGCs from literature (Supplemental Table 1) using hmmsearch (HMMER3). Rough bitscore significance cutoffs were determined for each pHMM. More precise cutoffs were assigned by testing against 28,688 NRPS BGC regions from the antiSMASH database (v3).^66^ BGC regions containing genes near the rough cutoff were manually inspected to determine if these were likely metallophore BGCs. If no clear bitscore cutoff could be discerned, representative low-scoring putative true hits were added to the pHMM seed alignment. This process was repeated until a precise bitscore cutoff could be determined.

### Phylogenetic analysis of Asp and His β-hydroxylases

Adequately-performing pHMMs for Asp and His β-hydroxylase subtypes could not be constructed using the above method. Siderophore β-hydroxylase functional subtypes were previously shown to form distinct phylogenetic clades.^26^ An expanded phylogenetic analysis was performed to serve as a guide for pHMM construction (Supplemental Dataset, `1_development/hydroxylase_tree/`). NRPS BGC regions from the antiSMASH database (v3) were scanned for matches to previously reported β-hydroxylase pHMMs^26^ and Pfam pHMMs for siderophore-related transporters (PF00593, PF01032, and PF01497^41,67^) using a modified version of antiSMASH v6.0. β-Hydroxylase genes meeting a relaxed bitscore cutoff of 300 (1070 total) were dereplicated with CD-HIT web server^68^ and a sequence identity cutoff of 70%, giving 425 representative amino acid sequences. A multiple sequence alignment was created using hmmalign (HMMER3) and the TauD Pfam (PF02668),^67^ and a maximum-likelihood phylogenetic tree was reconstructed with IQ-TREE (multicore v2.2.0-beta)^69^ using the WAG+F+I+G4 evolutionary model. The presence of nearby transporters was mapped onto the phylogenetic tree to identify clades or paraphyletic groups putatively involved in siderophore biosynthesis. Sequences in groups corresponding to previously reported TBH_Asp, IBH_Asp, and IBH_His subtypes and the novel putative CyanoBH_Asp1 and CyanoBH_Asp2 subtypes were extracted, and pHMMs were constructed and tested as described above.

### Incorporation into antiSMASH

The pHMMs and cutoffs were added to antiSMASH as a single detection rule called “NRP-metallophore” with the following logic:

~~~
VibH_like or Cy_tandem or
(cds(Condensation and AMP-binding) and (
        (IBH_Asp and not SBH_Asp) or IBH_His or TBH_Asp or
        CyanoBH_Asp1 or CyanoBH_Asp2 or
        IPL or SalSyn or (EntA and EntC) or
        (GrbD and GrbE) or (FbnL and FbnM) or PvdO or PvdP or
        (Orn_monoox and not (KtzT or MetRS-like))
        Lys_monoox or VbsL))
~~~

### Manual validation

RefSeq representative bacterial genomes were dereplicated at the genus level using R, randomly selecting one genome for each of the 330 genera determined by GTDB (Supplemental Dataset, `3_gtdb_taxonomy/` and `4_manual_testing/`). All NRPS BGC regions in the genomes were annotated with antiSMASH v6.1, yielding 758 BGC regions in the final testing dataset (Supplemental Table 2). The antiSMASH output for each BGC was manually inspected for evidence of NRP metallophore production, including genes encoding transporters, iron reductases, chelator biosynthesis, and known metallophore NRPS domain motifs. The same 758 BGC regions were classified as NRP metallophores using the chelator-based strategy described above, as well as the transporter-based strategy of Crits-Christoph *et al*.^*41*^ Genes matching Pfam pHMMs for siderophore-related transporters (PF00593, PF01032, and PF01497^41,67^) were identified using a modified version of antiSMASH v6.1, and BGC regions were classified as metallophores if two of the three transporter families were present.^41^ Each putative false positive was re-investigated before performance statistics were calculated, resulting in the reannotation of four BGCs.

### BIG-SCAPE clustering

NRP metallophore BGC regions from RefSeq representative genomes (Supplemental Dataset, `2_refseq_reps_results/metallophores_Jun25.tar.gz`) were filtered to remove clusters on contig edges. The resulting 2,523 BGC regions, as well as 78 previously reported BGCs (Supplemental Table 2) were clustered using BiG-SCAPE v1.1.2 with the following settings: “`--no_classify --mix --cutoffs 0.3 0.4 0.5 --clans-off`”. The network (Supplemental Dataset, `6_bigscape/mix_c0.50.network`) was imported to Cytoscape for figure preparation.

### Oxygen tolerance

Using the BacDive website,^54^ oxygen tolerance data, where available, were obtained for all bacterial strains that were associated with an NCBI genome, and filtered to include only strains with RefSeq representative genomes (resulting in 863 strains). Using R (Supplemental Dataset, `7_by_strain/bacdive_oxygen.rmd`) Strain oxygen tolerance data were summarized as *aerobe, anaerobe*, and *other* using the following rules: (1) “aerobe” or “obligate aerobe” were coded as *aerobe*, (2) “anaerobe” or “obligate anaerobe” were coded as *anaerobe*, (3) “facultative anaerobe”, “facultative aerobe”, or “microaerophile” were categorized as *other*, and (4) strains listed as any combination of *aerobe, anaerobe*, and *other* were coded as *other*.

### Phylogenetic mapping

A phylogenetic tree of 65,703 bacterial genomes was retrieved from the Genome Taxonomy Database (GTDB, version r207).^59^ Separately, RefSeq representative genomes were filtered to include only complete-level sequences. The GTDB tree was pruned using python to include only the intersection between both sets, resulting in 2,195 taxa (Supplementary Dataset, `3_gtdb_taxonomy/prune_tree.py`). Taxonomic lineages were also retrieved from GTDB, and suffixed taxa were combined (ex: “Firmicutes_A” was included in “Firmicutes”). Tree annotations were prepared using R scripts (Supplementary Dataset), and mapped using the interactive Tree of Life webtool (iTOL).^60^

## Data availability statement

All python, R, and bash scripts used in this paper, as well as underlying data, is available in the Supplemental Dataset, published to Zenodo: 10.5281/zenodo.7439477.

## Acknowledgements

Support from a European Research Council Starting Grant (948770-DECIPHER; ZR and MM) and the US National Science Foundation (CHE-2108596; AB) are gratefully acknowledged.

