## Supplemental Figures for "Automated genome mining predicts combinatorial diversity and taxonomic distribution of peptide metallophore structures"

**Figure S1**. Workflow for developing an NRP-metallophore-specific profile hidden Markov model (pHMM) and significance score cutoff for an enzyme (sub-)family. (1) Examples are collected from literature, the amino acid sequences are aligned with MUSCLE and (2) a pHMM is constructed with HMMER3. (3) BGCs of known function from MIBiG are scanned for matches to the pHMM to generate a preliminary bitscore cutoff. (4) NRPS BGC regions from the antiSMASH database are scanned for matches to the pHMM and sorted by bitscore. (5) Starting at the bitscore cutoff, BGCs are manually annotated to predict if they encode the biosynthesis of NRP metallophores using features such as genes encoding membrane transport, metal acquisition (ex: ferric reductases), and the biosynthesis of multiple chelating groups. In a properly functioning system, the bitscore cutoff delineates putative true and false positives to accurately detect the NRP-metallophore-related enzymes (bottom right). However, a bitscore cutoff adjustment may be required (bottom middle), or low-scoring true positives may need to be added to the pHMM seed alignment in an iterative process (bottom left).

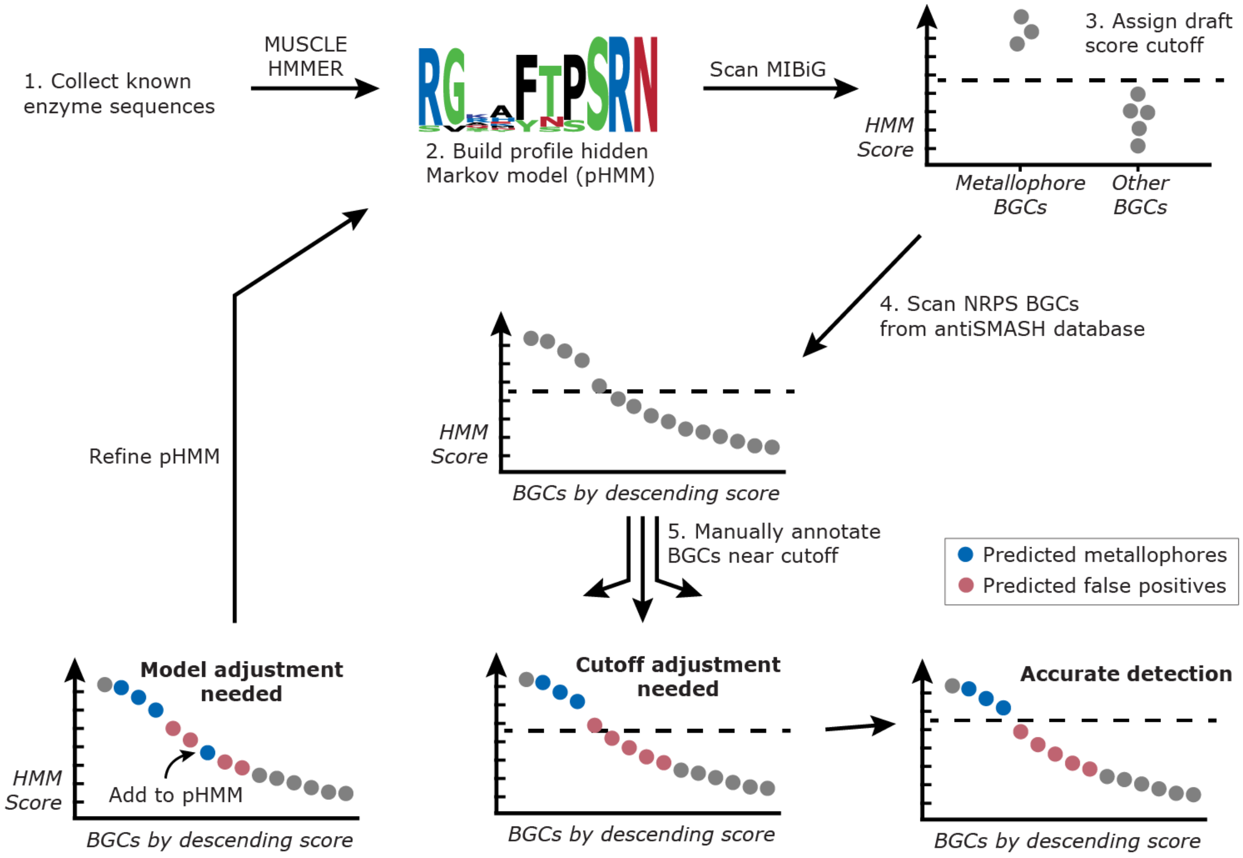

**Figure S2**. A maximum-likelihood phylogeny of putative β-hydroxylases found in NRPS BGC regions. β-Hydroxylase subtypes found in characterized siderophore BGCs are highlighted and labeled; the red SBH_Asp clade consists of non-metallophore phytotoxins.[1] Gray clades indicate possible metallophore β-hydroxylases that currently have no experimentally characterized representative. Amino acid sequences similar to known siderophore β-hydroxylase subtypes were extracted from the antiSMASH database (v3) and dereplicated prior to tree reconstruction. The right-hand bar gives the number of unique siderophore-related transporter families[2] present in the BGC region containing the β-hydroxylase.

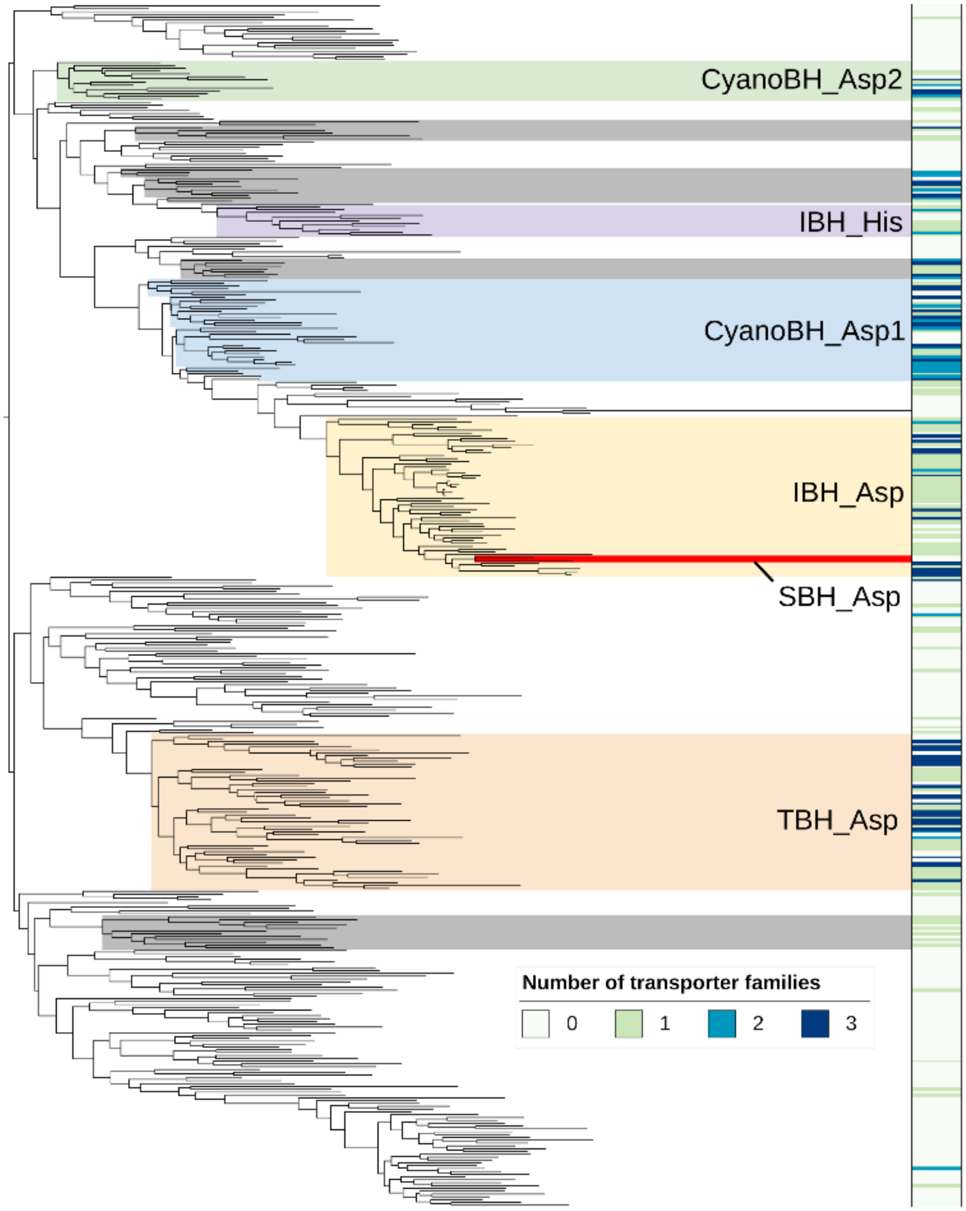

**Figure S3** (Next page). Analysis of a novel putative NRP metallophore BGC from *Sporomusa termitida* DSM 4440. (A) Clinker comparison of the *S. termitida* BGC with homologous loci. The *Sporomusa* sp. KB1 BGC contains a salicylate synthase gene detectable by the new antiSMASH rules, while *S. termitida* DSM 4440 instead contains several genes homologous to the menaquinone locus of *Desulfitobacterium* spp.[3] The cluster comparison was generated with clinker v0.0.26.[4] (B) A proposed metallophore biosynthesis pathway encoded by the *S. termitida* DSM 4440 BGC. 1,4-Dihydroxy-2-naphthoic acid is synthesized from chorismic acid by homologs of MenFDHBE, encoded by SPTER_RS05985-06005, and MenC, encoded elsewhere in the genome (SPTER_RS21050). The C4 phenol is likely methylated by O-methyltransferase SPTER_RS06015. The naphthoic acid moiety of Karamomycin C (inset box), characterized from an unsequenced strain of *Nonomuraea endophytica*,[5] is predicted to be synthesized by a similar pathway. Five NRPS genes in *S. termitida* DSM 4440 encode for the biosynthesis of the core structure by the condensation and cyclization of four Cys residues. The C-methyltransferase of SPTER_RS06025 is predicted to be inactive, as observed in the homologous ulbactin pathway.[6] The completed scaffold is released by thioesterase SPTER_RS05970, possibly producing the same tricyclic substructure observed in karamomycin C and ulbactin F (inset box). The final predicted structure accounts for the actions of two thiazoline reductases (SPTER_RS05950 and SPTER_RS05980) and a methyltransferase (SPTER_RS05955), although the the regiochemistry and timing of these transformations is unclear.

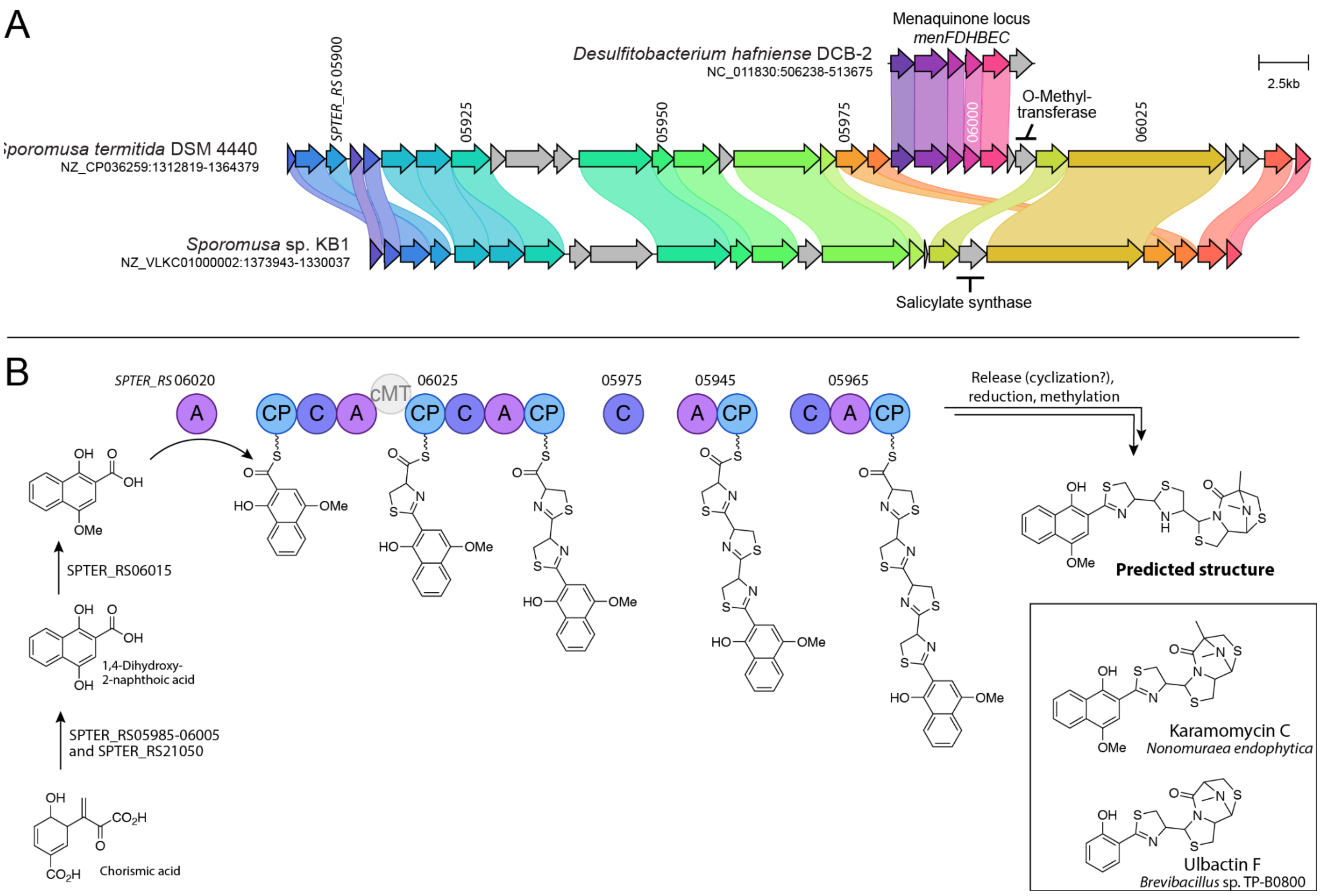
A
Sporomusa termitida DSM 4440
NZ_CP036259:1312819-1364379
SPTER_RS 05900
05925
05950
05975
06000
06025
Desulfitobacterium hafniense DCB-2
NC_011830:506238-513675
Menaquinone locus
menFDHBEC
O-Methyl-
transferase
2.5kb
Sporomusa sp. KB1
NZ_VLKC01000002:1373943-1330037
Salicylate synthase
B
SPTER_RS 06020
06025
05975
05945
05965
cMT
Release (cyclization?),
reduction, methylation
Predicted structure
SPTER_RS06015
1,4-Dihydroxy-
2-naphthoic acid
SPTER_RS05985-06005
and SPTER_RS21050
Chorismic acid
Karamomycin C
Nonomuraea endophytica
Ulbactin F
Brevibacillus sp. TP-B0800

**Figure S4**. Predicted structures of novel hydroxamate / β-OHHis metallophores from *Pseudomonas* spp.

Predicted siderophore from
*Pseudomonas xanthosomae* and *P. plecoglossicida*

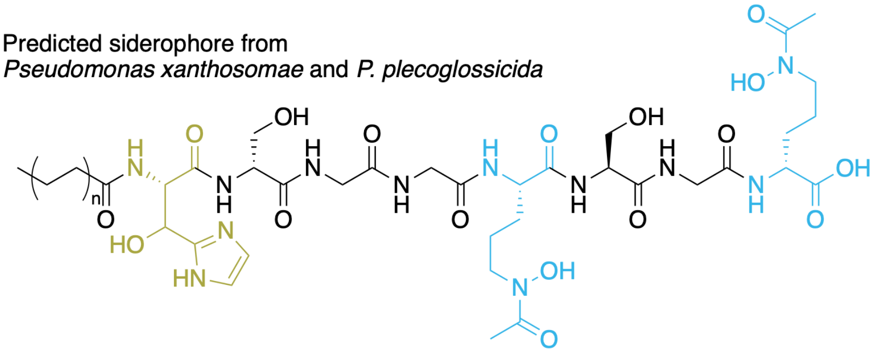

Predicted siderophore from
*Pseudomonas jinjuensis*

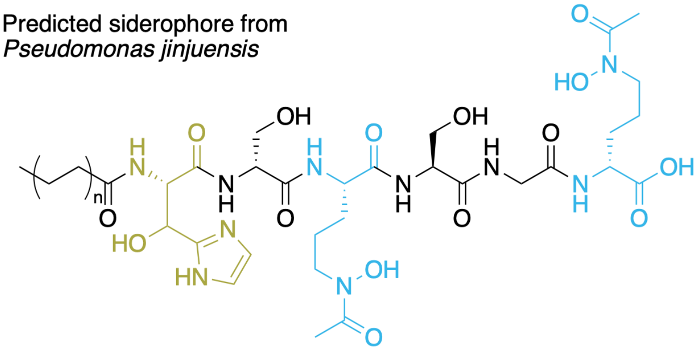

**Figure S5**. Similarity network of complete NRP metallophore BGC regions from RefSeq representative genomes. Nodes are colored blue if they belong to a GCF with a reference BGC, and orange if they are dissimilar from any reference BGC. The BiG-SCAPE network is identical to that in Figure 3 and reference BGC numbering corresponds to Supplemental Table 1.

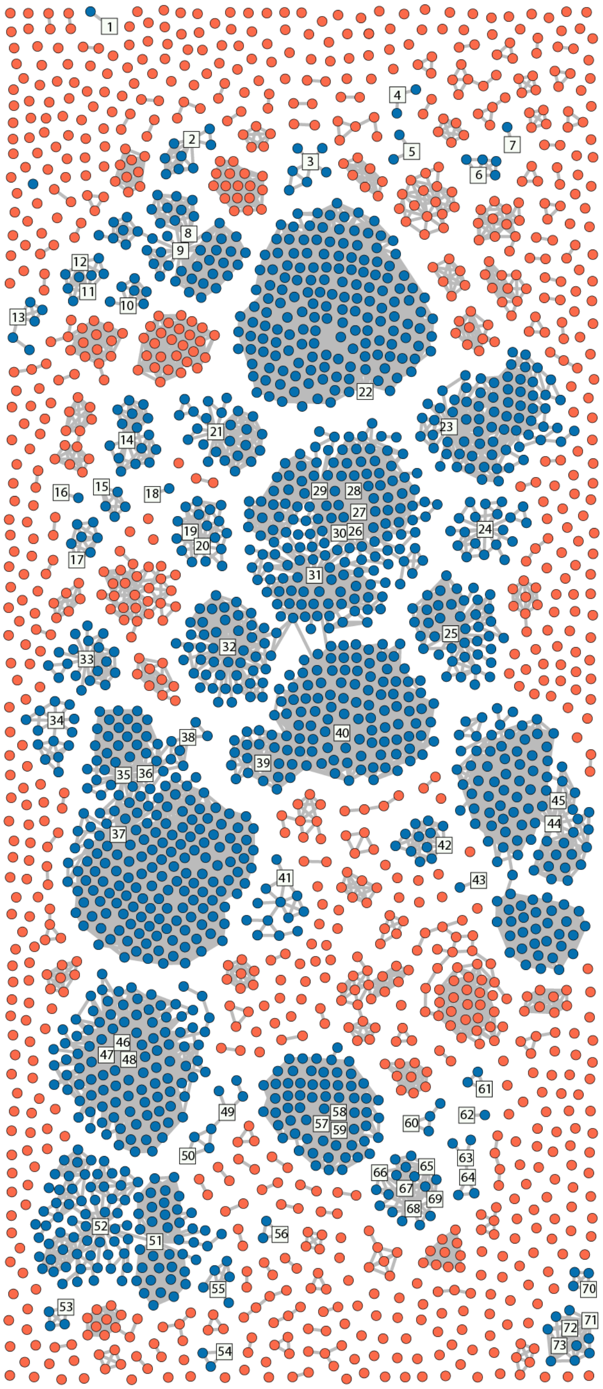

**Figure S6**. Taxonomic distribution of NRP metallophore BGCs (blue) and other NRPS BGCs (gray). Detected BGC counts per genome were mapped to the GTDB bacterial species tree with iTOL.

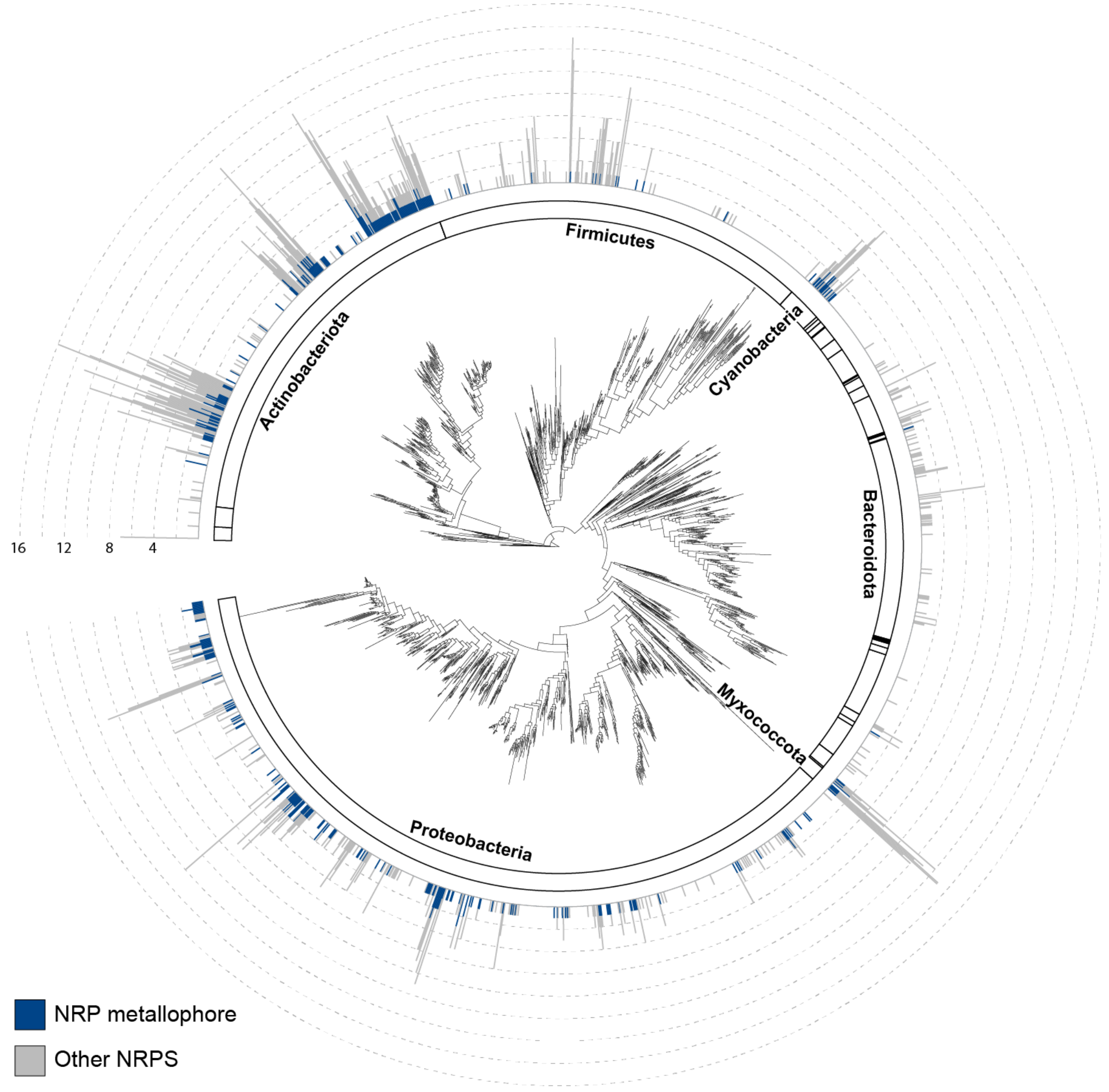

**Figure S7**. Taxonomic distribution of NRP metallophore BGCs similar (blue) and dissimilar (orange) from known reference BGCs. Detectable BGCs were mapped to the GTDB bacterial species tree with iTOL (identically to Figure 5, ring B) and classified according to whether they formed a gene cluster family with a known BGC (Figure S5).

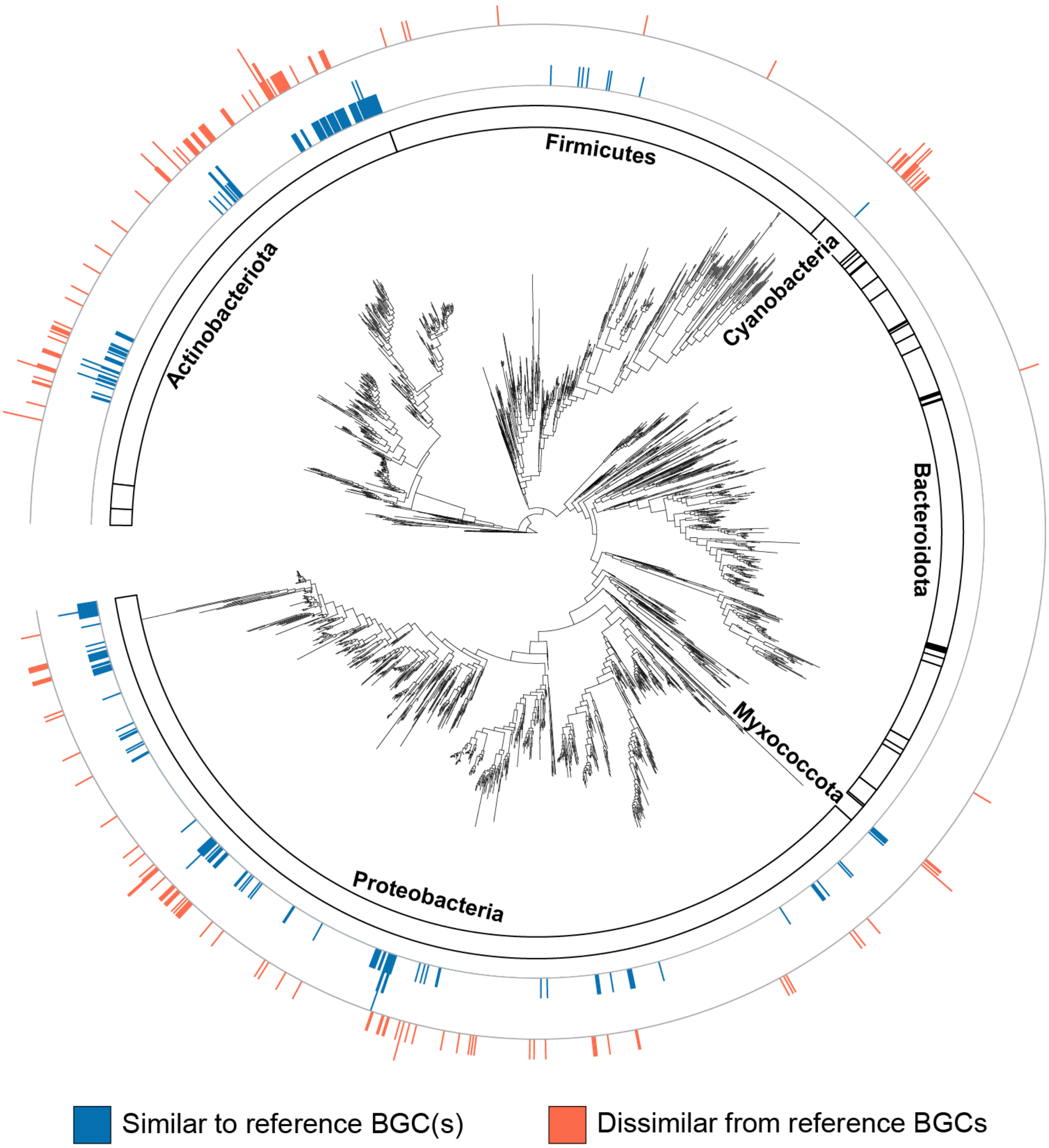

**Figure S8**. Taxonomic distribution of detected matches to individual metallophore pHMMs. Using iTOL, the GTDB bacterial species tree was annotated according to the presence of NRP metallophore BGCs containing (A) β-OHAsp pathways, (B) hydroxamate pathways, (C) salicylate pathways, and (D) metallophore-specific condensation domains.

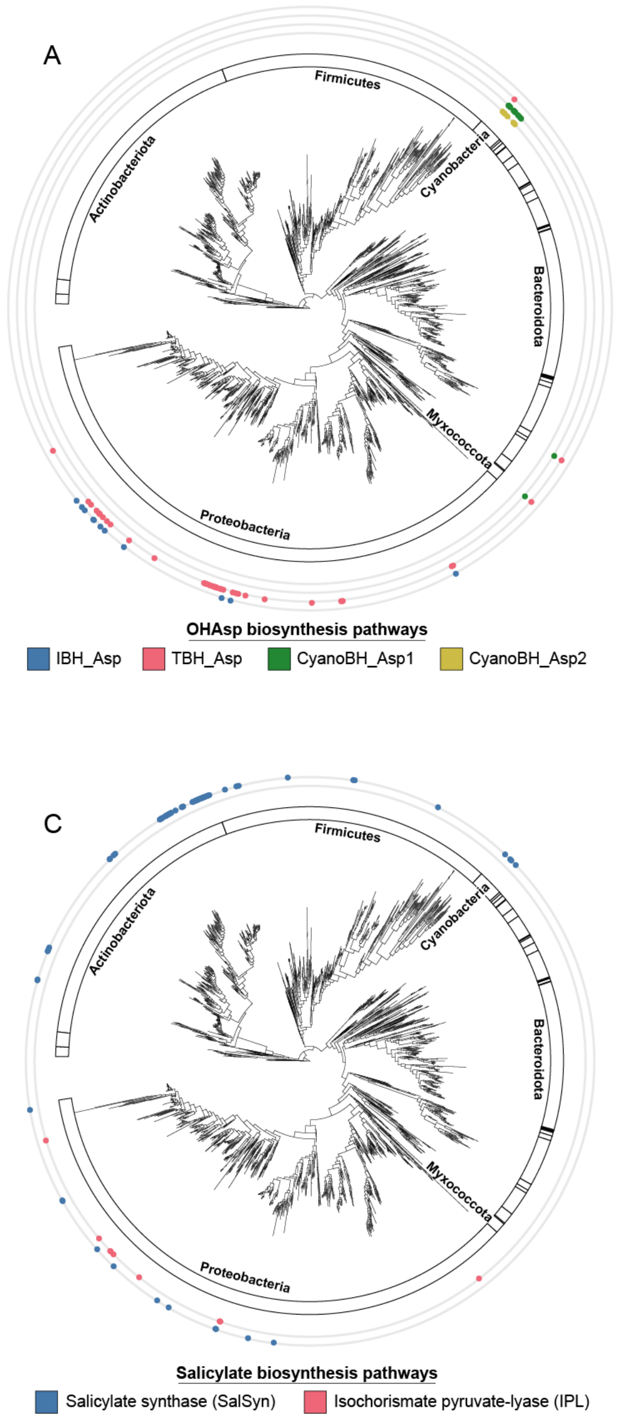
